# p53 mutations exhibit sex specific gain-of-function activity in gliomagenesis

**DOI:** 10.1101/2021.06.11.448124

**Authors:** Nathan Rockwell, Wei Yang, Nicole Warrington, Malachi Griffith, Obi Griffith, Christina Gurnett, Barak Cohen, Dustin Baldridge, Joshua Rubin

## Abstract

The tumor suppressor TP53 is the most frequently mutated gene in cancer. Most TP53 mutations are missense mutations in the DNA-binding domain, which in addition to loss of canonical p53 activity, frequently confer gain-of-function (GOF) aberrant transcriptional activity through mutant p53 localization to non-canonical genes. GOF phenotypes differ by mutation and cell identity and are reported to include increased proliferation, migration, metabolic reprogramming, and therapy resistance. We found that several recurring p53 mutations exhibit a sex-bias in patients with glioblastoma (GBM). *In vitro* and *in vivo* analysis of three mutations, p53^R172H^, p53^Y202C^, and p53^Y217C^ revealed sex differences in each mutation’s ability to transform primary mouse astrocytes. p53^R172H^ exhibited a far greater ability to transform female astrocytes than males, p53^Y202C^ transformed both male and female astrocytes with a small male bias, and p53^Y217C^ only exhibited GOF transformation effects in male astrocytes. These phenotypic differences reflect an interaction between sex and GOF mutation to drive unique gene expression patterns in cancer pathways. We found that mutant p53 exhibits sex and mutation specific aberrant genomic localization to the transcriptional start sites of upregulated genes, whose promoter regions were enriched for different sets of transcription factor DNA-binding motifs. Together, our data establish a novel paradigm for sex specific mutant p53 GOF activity in GBM with implications for all cancer.

## Introduction

The tumor suppressor *TP53* (p53), which regulates cell cycle progression, DNA-damage repair, apoptosis, and stem cell differentiation^1–5^, is the most commonly mutated gene in cancer^6,7^. It functions through direct protein-protein interactions and acting as a positive or negative regulator of transcription^8–10^ through its canonical DNA-binding site ^8–10^. p53’s transcriptional activity is dependent on many interacting and context specific factors such as the tissue or cell type, the mechanism of p53 activation, and the severity and longevity of DNA damage^11,12^. A growing body of evidence indicates that sex has a profound influence on the role of p53 in development, disease, and aging. In mice and rats, loss of p53 *in utero* results in decreased female progeny compared to males, as a consequence of aberrant X-inactivation in females, alters X-gene dosage, and results in embryonically lethal neural tube defects^13–15^.

p53 also exhibits sex differences in the maintenance of neural progenitor cells (NPC) in the sub-ventricular zone (SVZ)^16^. Under normal conditions, post pubertal male mice deplete NPCs faster than female. Knocking out p53 in these cells is protective against this depletion in males suggesting a male specific pro-apoptotic function of p53 in NPCs^16^. Regardless of geographical region, human females consistently exhibit greater longevity than males^17^. Work in drosophila has shown that tissue-general overexpression of WT p53 decreased lifespan in females and increased lifespan in males^18^. Tissue specific overexpression of WT p53 in the central nervous system had the opposite effect, increasing lifespan in females while decreasing lifespan in males^19^. The findings further support that p53 activity is both sex and tissue dependent.

Sex differences in the effects of p53 mutations have also been observed in cancer. Overall, in non-reproductive tumors, p53 mutations occur more frequently in males, and are associated with worse survival^20^. In individuals with Li-Fraumeni syndrome (LFS), a familial cancer predisposition syndrome associated with germline pathogenic variants in p53, females have an overall increased risk of developing cancer compared to males^21–25^, due to the large number of breast cancer cases. For some other cancers, such as brain tumors, males with LFS exhibit higher risk. Mice harboring a germline mutant p53 and loss of the promyelocytic leukemia (PML) gene, exhibit sex differences in cancer type and survival. Male mice develop more soft tissue sarcomas and have shortened survival compared to females, who are more likely to develop osteosarcomas and exhibit longer survival^26^. In a model of glioma, we previously observed that co-deletion of p53 and the tumor suppressor neurofibromin 1 (*Nf1*) is sufficient for transformation of male, but not female astrocytes^27^. Knocking out *Nf1* and p53 in NPCs *in utero* also leads to faster glioma formation and disease progression in male mice than female mice^28^.

The most common p53 alterations in cancer are missense mutations in the DNA-binding domain^29,30^. In addition to the loss of canonical DNA binding and gene regulation, many p53 missense mutations exhibit gain-of-function (GOF) and are more oncogenic than *TP53* deletion^31,32^. Many of these neomorphic functions can be attributed to p53-DNA interactions at non-canonical binding sites that drive the aberrant expression of oncogenes and repression of tumor suppressors^33,34^. The specific activity of different p53 mutations has been shown to be mediated by novel interactions with other proteins (such as NF-kB, ETS2, and NF-Y) and differential genomic localization^35–37^. Despite efforts to specify GOF mutant p53 phenotypes, characterizing the effects of mutant p53 GOF on transcription and tumorigenicity have been complicated by disparities in reported results, suggesting that, much like WTp53, mutant p53 activity, even the same missense mutation, is diverse and context dependent.

Given the known sex differences in p53 function and cancer incidence and survival, it is important to consider whether p53 GOF mutations contribute to sex differences in cancer^38^. Many of the pathways that contribute to sex differences in cell and systems biology including metabolism, cell cycle progression, invasion and EMT, epigenetic dysregulation, and immunity^38,39^, are associated with mutant p53 GOF activity. However, the influence of sex on mutant p53 GOF activity remains unknown. In glioblastoma (GBM), males have been shown to have both increased incidence and decreased overall survival compared to females^40^. Notably, *TP53* is among the most mutated genes in GBM with most mutations occurring as missense mutations in the DNA-binding domain^41^. Therefore, the intersection of sex differences in p53 function, sex difference in GBM incidence and outcome, and the high rate of p53 missense mutations in GBM make this an ideal system for interrogating the effects of sex on p53 gain-of-function activity with potential applicability to p53 GOF mutations in all cancer.

## Results

Across cancer, *TP53* mutations are most commonly missense mutations in the DNA binding domain (DBD) compared to deletions, inactivating (truncating mutations), or missense mutations outside of the DBD. In patients with Li Fraumeni Syndrome (LFS), central nervous system (CNS) tumors in particular are significantly associated with germline mutations in the helix-loop-helix region of the DBD, suggesting that these mutations preferentially predispose LFS patients to brain tumors^42^. To further investigate the association between p53 in CNS tumors and glioblastoma (GBM), we asked whether it was also present in somatic p53 mutations. *TP53* mutation data was aggregated from multiple studies deposited in the cBioPortal database, including the pan-cancer MSK-IMPACT Clinical Sequencing Cohort and all available studies for glioblastoma, pan-CNS cancers, and the top five cancer types by incidence: breast, lung, skin, pancreatic, and colorectal^43^. A complete list of studies and cancer subtypes included in this analysis can be found in **Supplementary Table S1**. We analyzed the frequency of missense mutations in the DBD as a percentage of all p53 mutations (**Figure 1A and 1B**). Despite the DBD contributing 48% of the amino acids in the full length p53 protein, missense mutations in the DBD account for 61.5% of p53 mutations in the pan-cancer dataset. In comparison to pan-cancer, missense mutations in the DBD are significantly more frequent in GBM (72.1%, p= 0.0001, Fisher’s Exact Test), CNS tumors (76.8%, p = <0.0001), and colon cancer (66.4%, p = <0.0001), significantly less frequent in breast cancer (56.5%, p =<0.0001), and skin cancer (52.5%, p = 0.0015), and equally common in prostate (61.4%) and lung cancers (61.7%). In total, missense mutations in the DBD were observed in 19.7% of GBM cases, compared to inactivating mutations in 7.6% and complete deletion in 1%. Together, these data support an enhanced impact of missense mutations in the p53 DBD on GBM and CNS tumor development.

**Figure 1.**
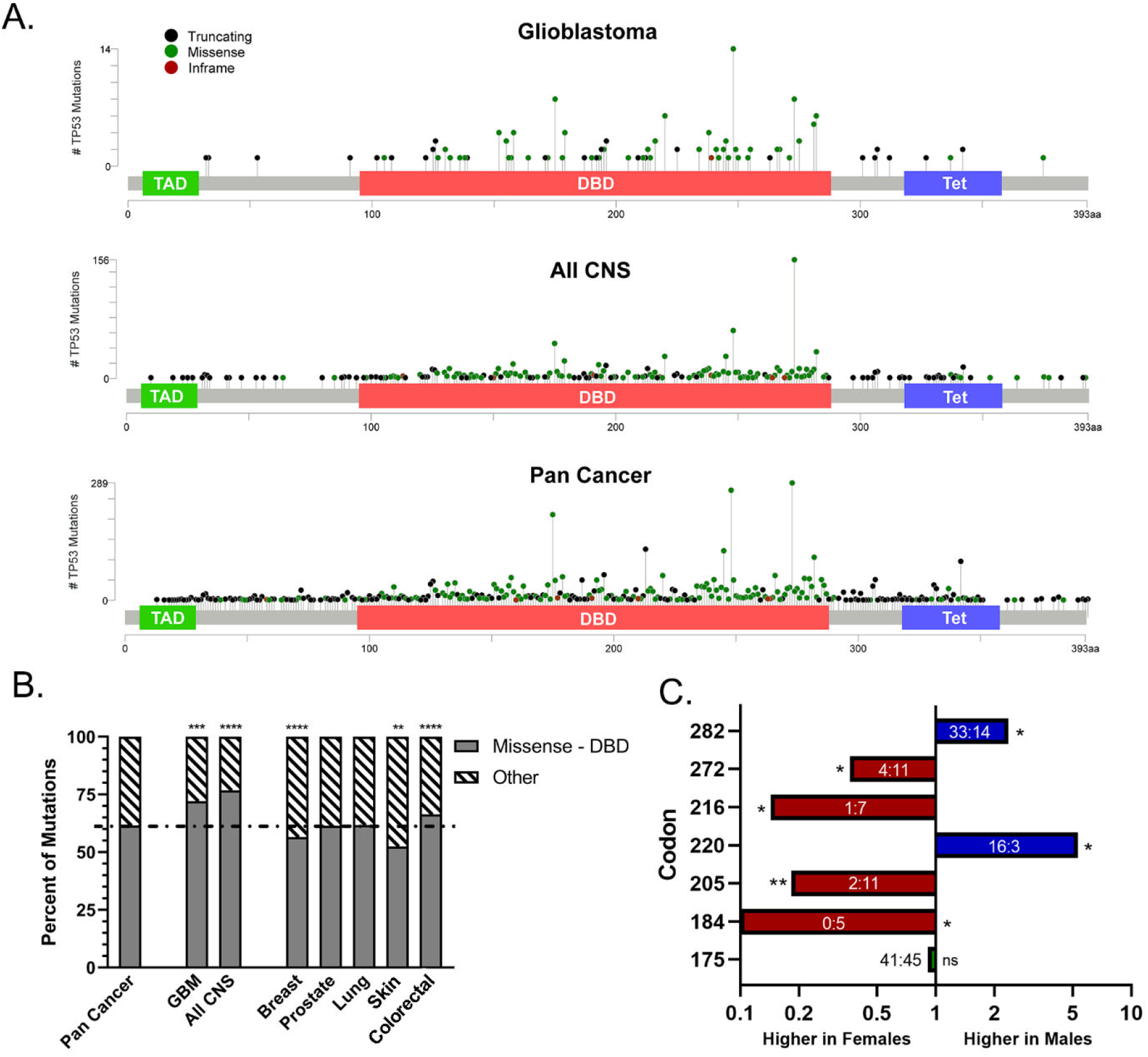
p53 missense mutations are enriched in GBM and show a sex difference in incidence. **A**. TP53 mutation analysis from cBioPortal representing distribution of all truncating, missense, and in-frame TP53 mutations in glioblastoma, all CNS tumors, and pan cancer analysis. **B**. The fraction of mutant TP53 tumors with missense mutations in the DNA-binding domain (Missense – DBD) was compared to all other TP53 mutation types and TP53 domains (Other) in pan cancer, GBM, CNS tumors, breast cancer, prostate cancer, lung cancer, skin cancer, and colorectal cancer. Significant enrichment or reduction compared to the pan cancer data set was determined by Fisher exact test. **C**. Analysis of the male to female ratio of TP53 missense mutations in GBM from TCGA, COSMIC, and IARC databases revealed six codons with a significant difference in the frequency of mutations in males and females. Bars are the ratio of male to female tumors with p53 mutated at each codon. The absolute mutation counts are labeled within each bar. Significance was determined by Fisher’s Exact test. * = p < 0.05, * * p = < 0.01, * * * = p < 0.001, * * * * = p < 0.0001

We next asked whether the frequency of these mutations is influenced by patient sex. *TP53* mutation data was combined and analyzed from GBM and CNS tumor specimens from The Cancer Genome Atlas (TCGA Cell 2013), the Catalogue of Somatic Mutations in Cancer (COSMIC), and the International Association for Research on Cancer (IARC) p53 database to determine the frequency of mutations at each codon in the DNA binding domain^6,44,45^. In total, 554 male tumors and 442 female tumors were included in the analysis. Six codons, present in 10.8% of all cases were identified as having a sex difference in their mutation frequency by Fisher’s Exact Test *p* < 0.05 **(Figure 1C)**. Four codons were mutated more frequently in females: D184, Y205, V216, and V272, and two codons more frequently in males: Y220 and R282. Notably, both male biased codons are classified as hotspot mutations for their high mutation frequency across cancers^46^. Codon R175 is among the best characterized hotspot mutations and has no sex difference in mutation frequency in GBM. The prevailing hypothesis for pan-cancer hotspot mutations is that these p53 GOF mutations confer a fitness advantage that drives their selection. Applying this same principle to our findings, we hypothesized that that p53 missense mutations have differential GOF effects in males and females that drive a sex biased selection of certain mutations.

### Modeling mutant p53 gain-of-function

To determine whether there are sex differences in p53 GOF activity, we developed a model of mutant p53 overexpression in mouse primary astrocytes. Briefly, astrocytes were harvested as previously described from the cortices of postnatal day one mouse pups harboring one p53 allele with flanking lox-P sites and one deleted p53 allele^27^. The pups were sex-typed using the X- and Y-chromosome paralogs, *Jarid1C/D*. Astrocytes from a minimum of three pups were combined per sex to produce male and female Trp53^f/-^ (p53 WT) astrocytes (**Figure 2A**)^47^. Astrocyte purity was assessed using glial fibrillary acidic protein (GFAP) immunofluorescence (IF) (**Figure 2B**). To evaluate potential sex-biased p53 GOF, we selected three mutations: *Trp53*^R172H^, *Trp53*^Y202C^, and *Trp53*^Y217C^ for evaluation. *Trp53*^R172H^ (Hs *TP53*^R175H^) is among the most common mutations across cancer and has established GOF^48^. It exhibits no sex difference in frequency. *Trp53*^Y202C^ (Hs *TP53*^Y205C^) is the mutation whose frequency is most significantly biased in female patients. *Trp53*^Y217C^ (Hs *TP53*^Y220C^) was the most significantly male biased mutation identified. Male and female p53 WT astrocytes were transduced with retrovirus to stably express mutant *Trp53*-IRES-*eGFP* for each missense mutation, and eGFP positive cells were sorted by flow cytometry. A common co-occurring event in cancers expressing mutant p53 is the loss of the second wild-type p53 allele, known as loss-of-heterozygosity (LOH). LOH has been shown to be required for stabilized mutant expression and GOF activity in some mutants^49^. To mirror these events in our *in vitro* cell model, WT and mutant p53 astrocytes were transduced with a lentiviral vector expressing Cre recombinase and Puromycin N-acetyl transferase (*Pac*). The cultures were selected for Cre/Pac positive cells for one week with 2.5 uM puromycin to produce p53 KO and p53^mutant/-^ astrocytes (**Figure 2C**). **Figure 2D** displays a representative western blot measuring p53 protein levels in each cell line. Under low stress conditions, WTp53 is negatively regulated by the E3 ubiquitin ligase Mdm2. In response to DNA damage p53 is phosphorylated inhibiting Mdm2 interactions and stabilizing the p53 protein. As a positive control, p53^f/-^ astrocytes were treated with a combination of the DNA-damaging agent etoposide (10 µg/ml) and the Mdm2 inhibitor Nutlin-3a (10 µM) or DMSO for eight hours leading to stable induction of p53 protein. p53 KO astrocytes expressing Cre but lacking mutant p53 expression had no detectable p53. In contrast, high levels of overexpressed mutant p53 protein were observed in male and female astrocytes transduced with retrovirus to express all three mutations. To confirm the loss of the WT floxed p53 allele in Cre-positive cells we performed a PCR with primers specific to the floxed and deleted p53 alleles (**Figure 2E**). Introduction of Cre eliminated the floxed allele band indicating the complete loss of p53 at the WT locus. To ensure that WTp53 was completely depleted in the Cre expressing astrocytes, WTp53 and p53KO astrocytes were treated with etoposide and Nutlin-3a or DMSO for 8 hours and then p53 protein detected by IF. Co-treatment induced high levels of nuclear localized p53 in WT astrocytes but resulted in no detectable p53 in the p53 KO astrocytes (**Figure 2F**). IF for p53 in male and female astrocytes expressing mutant p53 confirmed that in the absence of treatment all three mutations expressed high levels p53 protein throughout the cell body (**Figure 2G**).

**Figure 2.**
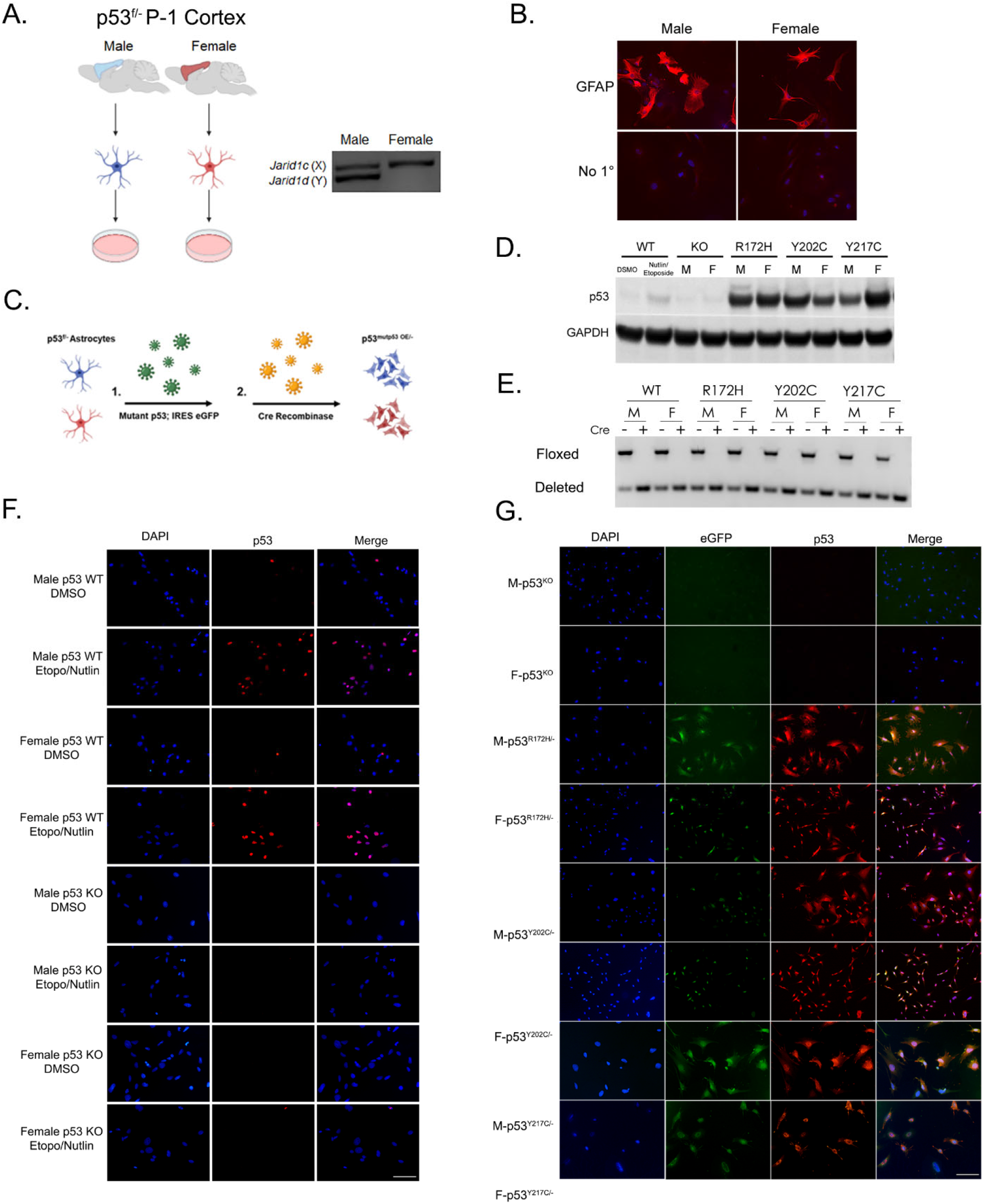
Modeling p53 missense mutations in GBM. **A**. Trp53^f/-^ astrocytes were isolated from the cortices of male and female postnatal day 1 mouse pups. The sex of the astrocytes was determined by genotyping PCR (left panel), and male or female astrocytes from at least 3 pups were combined. **B**. Astrocytes were stained for the astrocyte lineage marker GFAP to confirm astrocyte purity. Scale bar = 100 µm **C**. To create mutant p53 expressing astrocytes, Trp53^f/-^ astrocytes were transduced with retrovirus to overexpress mutant Trp53-IRES-eGFP, and eGFP positive cells were sorted by flow cytometry. Sorted eGFP positive astrocytes were transduced with lentivirus to express Cre-IRES-Puro and selected with 2.5 ug/ml puromycin for 1 week. **D**. Western blot assay was performed using 50 ug of whole cell lysates from male and female p53^f/-^ (WT), Trp53 KO, and Trp53^mut/-^ astrocytes. Trp53^f/-^ (WT) astrocytes were treated with 20 ug/ml etoposide and 10 ug/ml Nutlin-3a or DMSO control for 8 hours. Mutant p53 astrocytes stably express mutant p53 protein at high levels compared without the treatment needed to stabilize WT p53 protein. **D**. Loss of the lox-p53-lox (floxed) allele following introduction of Cre recombinase was confirmed by PCR. **F**. Treatment with Nutlin-3a and etoposide to stabilize p53 results in detectable p53 protein in p53 WT but not p53 KO astrocytes. Trp53^f/-^ astrocytes were treated with 20 ug/ml etoposide and 10 ug/ml Nutlin-3a or DMSO for 8 hours and were fixed and stained by immunofluorescence for p53 protein. Scale bar = 200 µm **G**. IF for GFP (green) and p53 (red) p53 KO and mutant astrocytes. Scale bar = 200 µm

### Sex differences in p53 GOF phenotype

Accelerated proliferation is among the most frequently observed phenotypes attributed to mutant p53 GOF. In addition to the loss of WTp53 regulation of cell cycle, GOF mutant p53 interacts with several transcription factors to increase the transcription of pro-proliferation genes^48^. Thus, we measured the growth effects of each mutation in both sexes. 3,000 male or female p53 WT, p53 KO, p53^R172H^, p53^Y202C^, and p53^Y217C^ astrocytes were plated in triplicate on 10 cm dishes, in three independent experiments, and allowed to expand for eight days before fixing and Giemsa staining nuclei. KO alone increased astrocyte proliferation compared to WT astrocytes but exhibited no detectable sex differences. All three p53 mutations exhibited a GOF growth phenotype in a sex dependent manner. Expression of p53^R172H^ in females and p53^Y202C^ and p53^Y217C^ in males significantly increased growth in comparison to p53 KO astrocytes and astrocytes expressing the same mutation in the opposite sex. While p53^R172H^ in males, p53^Y202C^ in females, and p53^Y217C^ females were insufficient to increase proliferation over p53 KO (**Figure 3A and C**), male p53^R172H^ and female p53^Y202C^ astrocytes formed denser colonies compared to KO, suggesting that these mutations may still possess sex-biased GOF activity (**Figure 3B**). In female cells, p53^Y217C^ expression appeared to decrease the growth phenotype compared to KO. To ensure that this effect was not a technical artifact of the assay, we plated a 5-fold titration of male and female p53 KO and p53^Y217C^ astrocytes and incubated them for five days at which point the fastest growing cells reached over- confluency (**Figure S1**). This experiment validated the previous observation, with female p53^Y217C^ growing slower than p53KO, p53KO exhibiting no sex difference, and male p53^Y217C^ growing the fastest. Together, these data support that p53 GOF effects are dependent on the interaction between sex and mutation.

**Figure 3.**
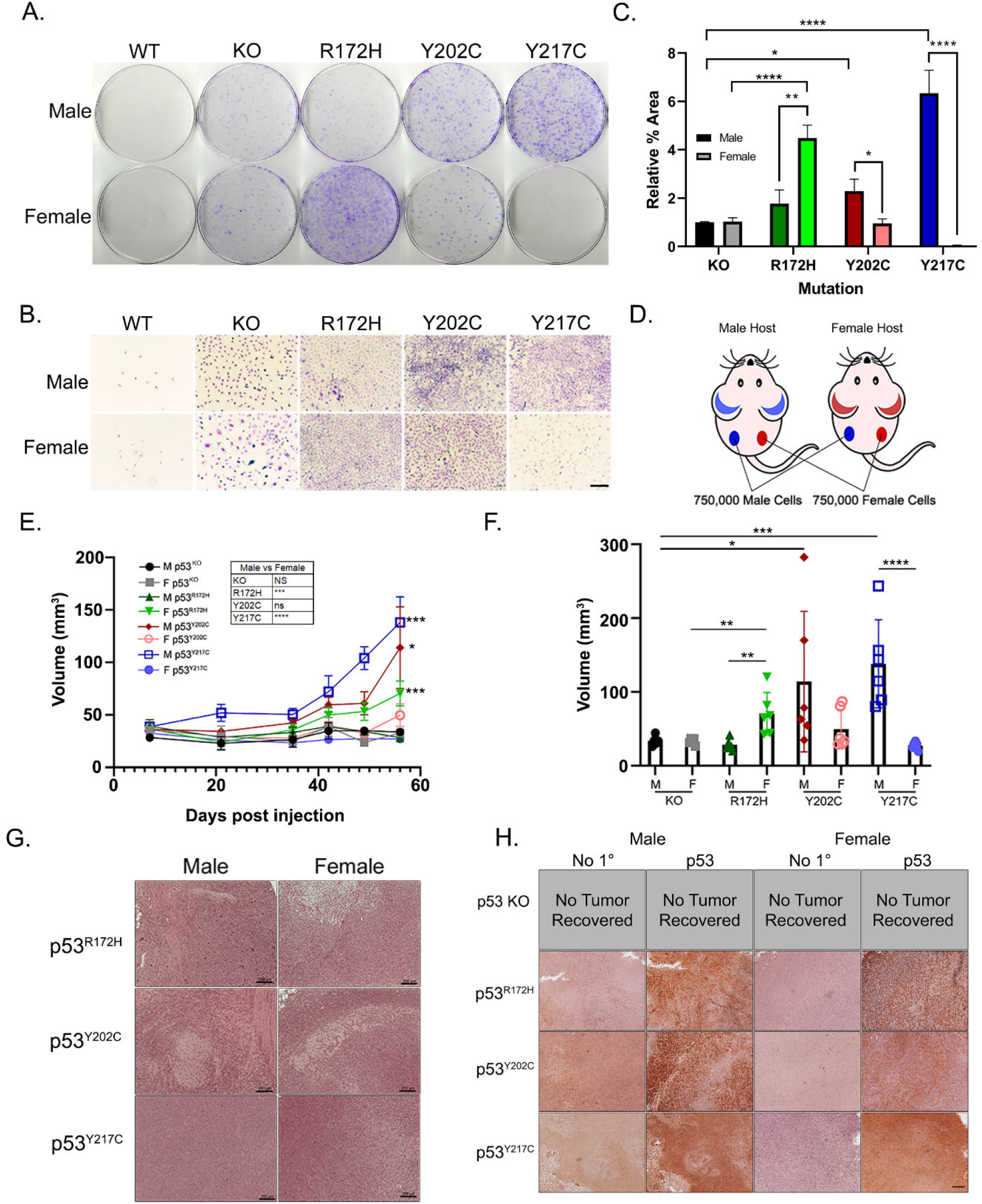
p53 missense mutations exhibit sex specific GOF effects on growth and tumorigenicity. **A**. Male and female WT p53, p53 KO, p53^R172H^, p53^Y202C^, and p53^Y217C^ astrocytes were plated at 3000 cells per 10 cm and incubated in normal growth conditions for 8 days, fixed with 100% methanol, and stained with Giemsa stain. Representative plates show a sex specific gain-of-function growth phenotype. **B**. Representative magnified images of astrocyte colonies showing differences in the density of colonies between male and female mutant p53 astrocytes. Scale bar = 100 µm. **C**. The percent area covered by stained cells was quantified in ImageJ and normalized to male KO controls. n=3 Significance determined by unpaired student’s T-test **D**. Schematic of subcutaneous flank injection protocol. 750,000 male and female astrocytes were injected subcutaneously into the opposing flanks of 3 male and 3 female nude mice. **E**. Flank tumors were measured weekly using a digital caliper. n=6. Female p53^R172H^, male p53^Y202C^, and male p53^Y217C^ tumors grew significantly faster than their sex matched p53 KO astrocytes. p53^R172H^ and p53^Y217C^ tumors also exhibited sex differences in tumor growth (inset table). Significance was determined by two-way ANOVA. **F**. Tumors were harvested from the mouse flanks and volumes measured post resection. When no tumor was recovered, the original Matrigel injection was measured for final tumor volume. Significant differences in tumor volume were determined by performing a log transformed Welch-corrected two-tailed t-test. **G**. Representative hematoxylin and eosin-stained sections from flank tumors. Scale bars = 200 µm. **H**. Immunohistochemical staining was performed on each resected tumor showing high expression of mutant p53 in the tumor. Scale bar = 200 µm. * = p < 0.05, * * = p < 0.01, * * * = p < 0.001, * * * = p < 0.0001.

Next, we asked whether the growth phenotypes were correlated with sex differences in tumorigenicity. Equal numbers of each male and female KO and p53 mutant astrocyte lines were injected into parallel flanks of three male and three female NCr nude mice (n=6) as illustrated in **Figure 3D**. Flank tumor volume was measured by digital caliper weekly until the first tumor reached terminal size. Male p53^Y217^, male p53^Y202C^, and female p53^R172H^ tumors grew significantly faster than p53 KO tumors (two-way ANOVA, *p <* 0.05) (**Figure 3E**). When comparing between sexes, male p53^Y217C^ and female p53^R172H^ tumors grew significantly faster than the same mutation in the opposite sex, while neither p53^Y202C^ nor p53 KO exerted sex-biased effects (**Figure 3E inset**). In total, 0/6 KO, 16.7% (1/6) of male and 83.3% (5/6) of female p53^R172H^, 66.6% (4/6) of male and 50% (3/6) of female p53^Y202C^, and 100% (6/6) of male and 16.6% (1/6) of female p53^Y217C^ astrocyte injections formed tumors. In addition to the frequency of tumor formation, female p53^R172H^, male p53^Y202C^, and male p53^Y217C^ tumors were significantly larger than tumors expressing the same mutation in the opposite sex (**Figure 3F, Figure S2**). When no tumor was recovered, the final tumor volume reflects the volume of the recovered Matrigel pellet. Resected tumors all had features indicative of gliomas including nuclear atypia and hypercellularity (**Figure 3G**) and expressed high levels of mutant p53 protein (**Figure 3H**). In comparison to p53 KO, all mutations exhibited a GOF in tumorigenicity compared to p53 loss regardless of sex. However, three distinct patterns of sex-biased GOF were observed. p53^R172H^ mutants had the greatest female biased sex effect. p53^Y202C^ had a bias toward greater tumorigenicity in males but had the smallest differences between sexes. p53^Y217C^ GOF activity was greatest in males and was the only mutation which formed tumors from every injection supporting male specific GOF for this mutation.

### Mutant p53 and genomic instability

All mutations were introduced at the same astrocyte passage under equivalent conditions with WTp53 intact. Introduction of mutant p53 and loss of WT p53 function disrupts DNA repair and cell cycle checkpoints leading to genomic instability and the acquisition of additional mutations^50^. We questioned whether the observed sex differences in growth and tumorigenicity might be driven by differences in genomic instability or acquisition of sporadic oncogenic mutations. We performed whole exome sequencing on cell expressing all three mutations and p53 KO in males and females and measured mutational burden. Loss of WT p53 function led to equivalently large increases in mutational burden in all eight cell lines. To determine whether overall mutational burden was impacting phenotype, the raw mutation count was plotted against the percent area from the growth assay in Figure 2. There was no significant correlation between mutational burden and proliferation (**Figure S3A**). Most mutations identified were silent or intergenic. Further analysis of missense, frame-shift, and non-sense mutations predicted to have a moderate to high impact on gene expression or function, also exhibited similar frequencies between cell lines with no correlation between the number of mutations and growth (**Figure S3B**).

Finally, we asked whether there were any unique mutations that could be contributing to these phenotypic differences. **Table S2** lists every gene with a homozygous missense, frame-shift, or nonsense mutation in at least one cell line. Almost all mutated genes were altered across tumor forming and non-tumor forming cells. Three genes, Calcoco2, Rbmy, and Thap1 were identified as uniquely mutated only in male p53^Y202C^ cells. None of these genes have known functions associated with cancer related pathways. Together, these data suggest that genomic instability and the acquisition of sporadic mutations does not account for the observed sex differences.

### Sex and mutant p53 interact to drive unique transcriptional profiles in oncogenic pathways

One of the primary mechanisms of mutant p53 GOF activity is the transcriptional regulation of genes in pro-tumorigenic pathways. Previous studies have identified several oncogenic targets of both R175H and Y220C mutations^48^. To determine whether mutant p53 exhibits GOF transcriptional activity, we performed mRNA-sequencing (RNA-seq) and compared the differentially expressed genes between each mutation and p53 KO within each sex. In all cases, introduction of mutant p53 led to significant changes in gene expression compared to p53 KO, confirming neo-morphic transcriptional function (**Figure S4**). We next compared the differentially expressed genes between male and female astrocytes within each mutation. p53 KO, p53^R172H^, p53^Y202C^, and p53^Y217C^ yielded 7183, 9013, 10,902, and 10,748 significant differentially expressed genes between sexes, respectively (FDR < 0.05) (**Figure 4A**). Predictably, sex chromosome linked genes including the X-linked genes *Xist, Agtr2*, and *Eda*, and the Y-linked genes *Uty, Eif2s3y, Ddx3y*, and *Kdm5d* are among the most differentially expressed genes across all mutations. However, these genes represent only a small fraction of sex-specific gene expression.

**Figure 4.**
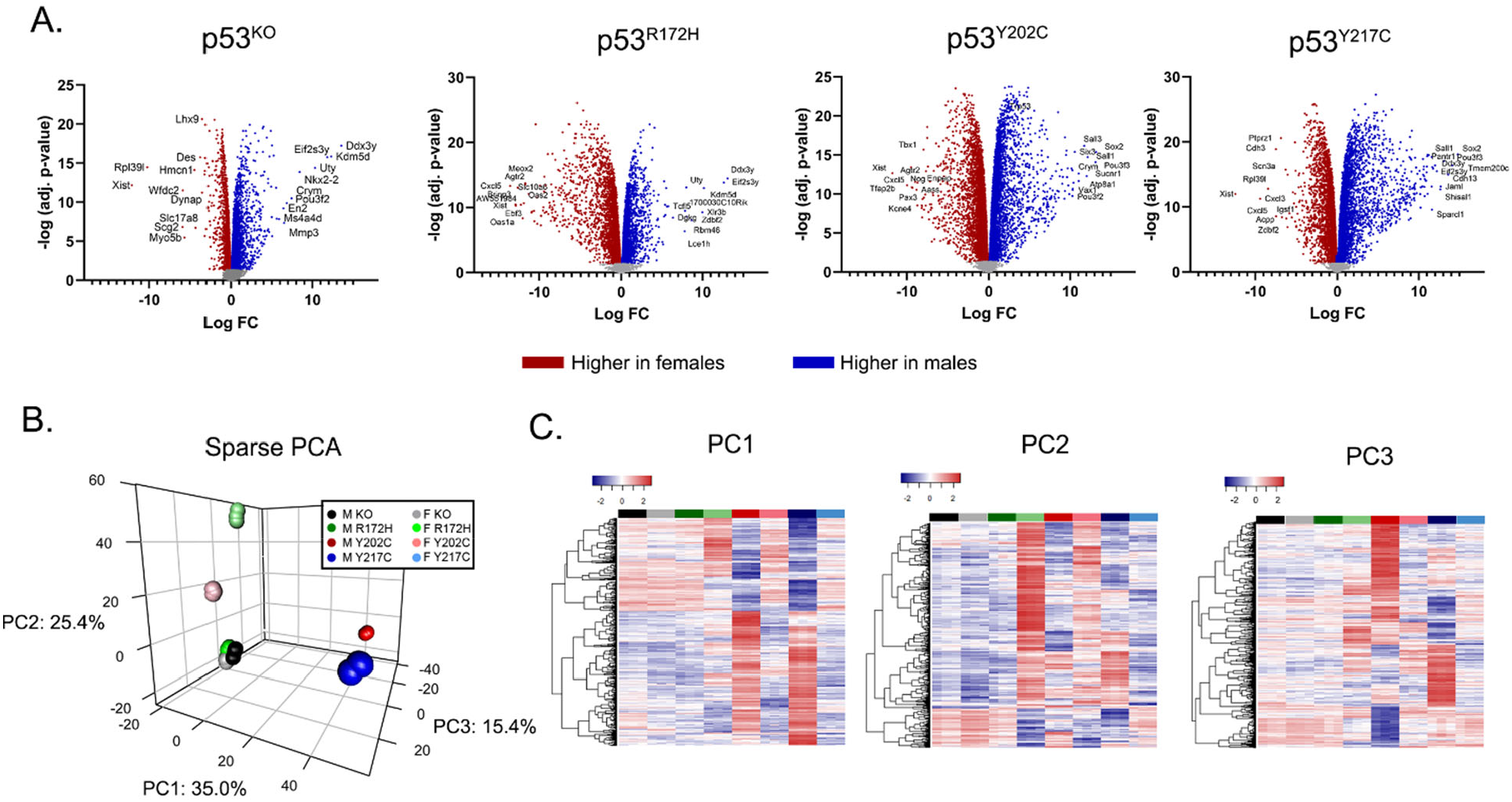
p53 missense mutations have sex specific/mutation specific GOF effects on gene expression. **A**. Volcano plots of all significant differentially expressed genes between males (blue) and females (red). The labelled points represent the ten genes with the greatest differences in expression (logFC). Significance was determined by FDR < 0.05. **B**. Sparse PCA analysis displaying variation between highly tumorigenic and weakly-tumorigenic cell lines. **C**. Heatmaps with hierarchical clustering of log2 transformed CPM for all genes contributing to each principal component in the sparse PCA analysis. Each colored bar spans three independent replicates for each mutation in each sex. Colors matched the figure legend in panel B.

To evaluate how gene expression varies as a function of p53 mutation and sex, we performed a sparse principal component analysis. The first three principal components account for 75.8% of total variance. Strikingly, the samples clustered primarily by their ability to form tumors with low-tumorigenic male and female KO, male p53^R172H^, and female p53^Y217C^ clustering together, while high-tumorigenic female p53^R172H^, female p53^Y202C^, male p53^Y202C^, and male p53^Y217C^ clustered independently (**Figure 4B**). Among the cell lines with high-tumorigenic potential, the first two principal components account for the majority of sex differences with male p53^Y202C^ and p53^Y217C^ separating in PC1 and female p53^Y202C^ and p53^R172H^ separating in PC2. Meanwhile, PC3 encompasses the majority of variation between mutants. The normalized expression for all genes contributing to each principal component are displayed in heatmaps in **Figure 4C**. All tumor-forming cell lines have distinct transcriptomes, suggesting that neither sex nor p53 mutation status are independently sufficient to explain variation in gene expression. Rather, it is the interaction between mutation and sex that drives unique transcriptional phenotypes.

To characterize the sex specific GOF transcriptome of each mutation, differentially expressed genes between mutant p53 and p53 KO were compared between sexes to define the sex specific (unique) and shared (overlapping) transcriptional impact of each mutation. We focused our analysis on those genes with an absolute log(fold change) change greater than 0.5. KEGG pathway enrichment analysis was used to evaluate whether shared or sex specific changes in gene expression could be contributing to the observed differences in tumorigenic potential. Concordant with differences in tumorigenesis, pathway analysis revealed three independent transcriptional patterns. p53^R172H^ expression had a substantially greater impact in females than males with approximately 7.4x more female specific differentially expressed genes (**Figure 5A**). Pathway analysis revealed that shared differentially expressed genes were enriched for KEGG cancer pathways, highlighted in orange. Additionally, shared genes were enriched for many cancer-associated pathways including cell cycle, DNA replication, p53 signaling, focal adhesion, senescence, and Rap1 signaling. The genes differentially expressed uniquely in males were not enriched for any cancer associated pathways. Meanwhile, genes that were differentially expressed in females alone were also enriched for many KEGG cancer pathways and cancer-associated pathways including ECM-receptor signaling, stem cell pluripotency, and PI3K-Akt signaling among others. This indicates that p53^R172H^ has both female specific and sex independent GOF effects on gene expression in cancer relevant pathways, and the combined effect is sufficient for tumorigenesis in females, while the universal GOF alone is insufficient in males.

**Figure 5.**
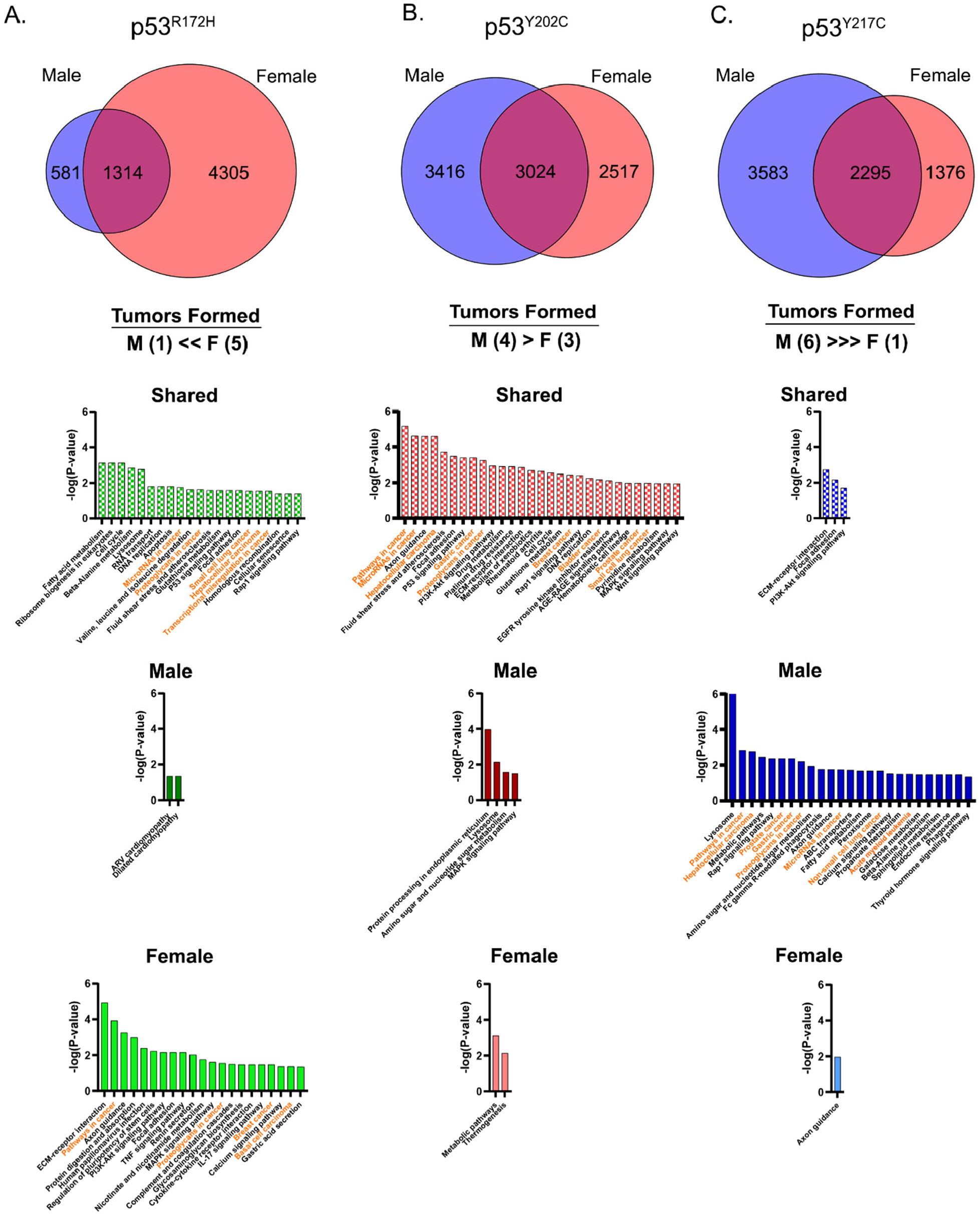
Sex and mutation interact to drive gene expression patterns in cancer pathways concordant with the tumorigenic phenotype. All differentially expressed genes (logFC > 0.5, FDR < 0.5) between each mutation and p53 KO were compared to define the sex specific and shared GOF transcriptional profile. Venn diagrams display the number of shared and sex specific differentially expressed genes. KEGG pathway enrichment analysis applied to each corresponding gene list revealed sex and mutation specific patterns of enrichment for changes in cancer associated pathways (orange) for p53^R172H^ (**A**), p53^Y202C^ (**B**), and p53^Y217C^ (**C**). The magnitude of sex difference in GOF phenotype (number of tumors formed *in* vivo) is displayed below each Venn diagram.

The p53^Y202C^ mutation had the smallest sex difference in GOF phenotype with both males and females forming large tumors *in vivo*, and the smallest differences in sex specific genes of any mutation. For this mutation, the overlapping gene list was enriched for cancer pathways and cancer-associated pathways with very few enriched pathways in the sex specific differentially expressed genes (**Figure 5B**). Together this reveals a second GOF transcriptional profile in which the universal GOF activity is sufficient for transformation in both sexes with very little sex specific effect on transformation.

The p53^Y217C^ mutation represents a third interaction between mutation and sex (**Figure 5C**). Here, male specific genes were the only subset enriched for KEGG cancer pathways, with very little pathway enrichment in the shared or female component. Male specific genes were enriched for pathways in cancer, Rap1 signaling, proteoglycans in cancer, microRNAs in cancer, several tissue specific cancer pathways, and cancer associated metabolic pathways. These data support a model in which p53^Y217C^ has male specific GOF effects on transcription with little universal or female specific activity in cancer relevant pathways.

### Sex differences in mutant p53 GOF activity is mediated through differential genomic localization

Having established that mutant p53 expression leads to unique GOF transcriptional profiles dependent on both mutation and sex, we next sought to determine the mechanism underlying the differences mutant p53 transcriptional regulation. We performed chromatin immunoprecipitation with massively parallel DNA sequencing (ChIP-seq) to map mutant p53 localization. To understand the sex differences in mutant p53 function, we focused on the mutation with the greatest female specific effect, p53^R172H^, and greatest male specific effect, p53^Y217C^. Regions enriched for mutant p53 localization were then mapped relative to their distance from the transcriptional start sites (TSSs) of the top 1000 upregulated genes for each mutation in males and female (**Figure 6A**). In both males and females, p53^R172H^ localization was enriched near the TSSs of a subset of upregulated genes indicating that mutant p53 directly regulates gene expression in both sexes, which supports GOF transcriptional activity in both sexes. In p53^Y217C^ cells, localization was enriched at TSSs of upregulated genes in males but displayed no pattern of localization in females.

**Figure 6.**
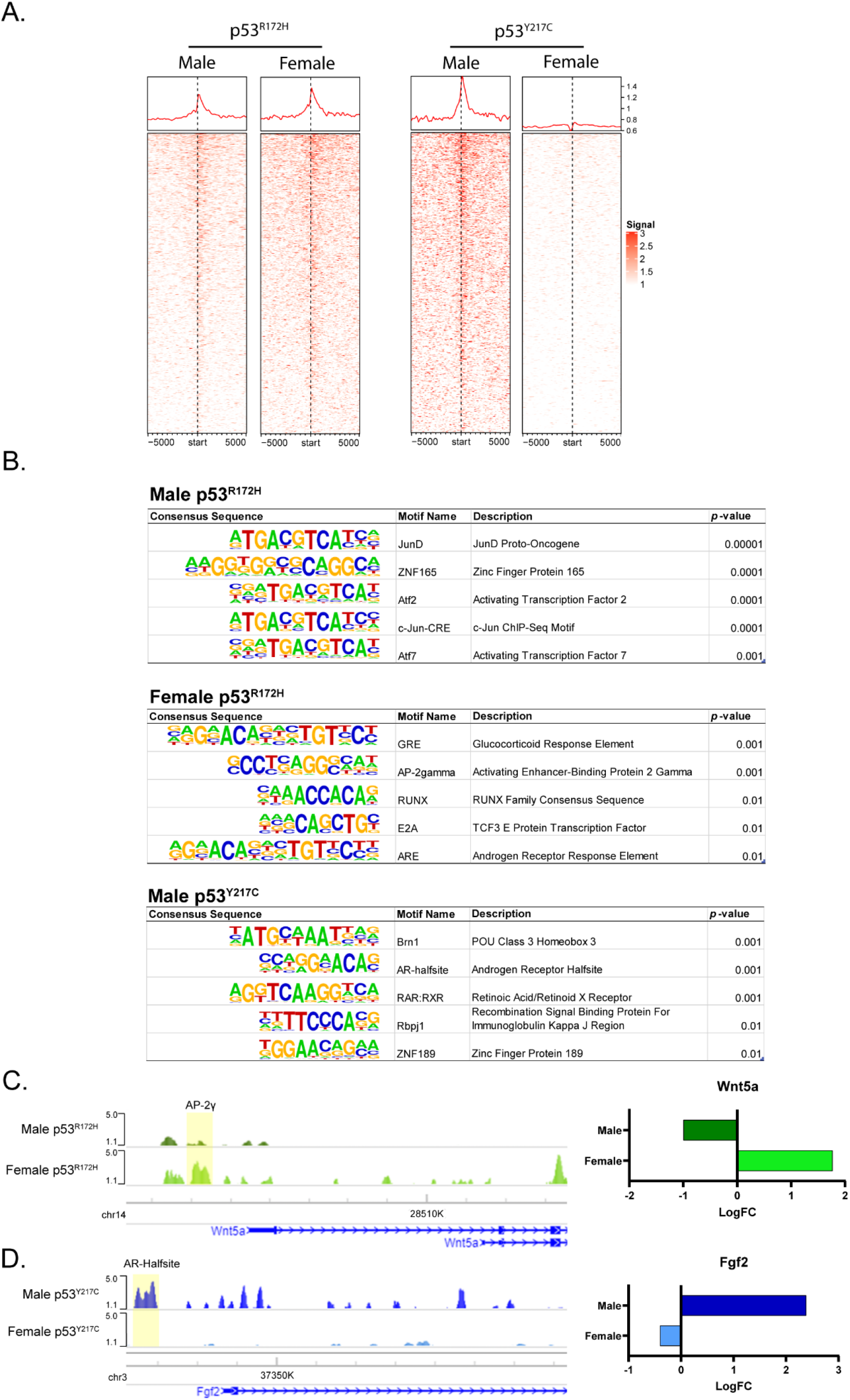
ChIP-seq revealed mutant p53 localization at transcriptional start sites is enriched for sex/mutation specific transcription factor DNA-binding motifs. **A**. Heatmaps of mutant p53 ChIP-seq enrichment over input +/- 5 kb from transcriptional start sites of the top 1000 upregulated genes compared to p53 KO for male and female p53^R172H^ and p53^Y217C^. The line graph above each heatmap displays the average signal across all 1000 genes. **B**. Homer analysis of p53^R172H^ in males and females and p53^Y217C^ in males within +/- 5 kb of the TSSs. The top five significantly enriched known consensus sequences for each mutation and sex are shown. **C**. Female enriched p53^R172H^ localization at the AP-2γ motif (yellow) in the TSS of *Wnt5a* correlates with increased gene expression compared to males. **D**. Male specific p53^Y217C^ localization at the AR-halfsite motif (yellow) in the TSS of *Fgf2* correlates with increased gene expression compared to females. Bar graphs display the logFC in gene expression between each mutation and p53 KO with the same sex.

Previous studies have established that mutant p53 lacks a consensus binding sequence, but instead relies on interactions with other transcription factors and DNA-binding proteins for localization to aberrant target genes. To identify potential mediators of mutant p53 activity, we performed a Homer analysis on the enriched ChIP signals +/- 5 kb of the TSS of the same 1000 upregulated genes enriched for p53 localization. The top five significant known motifs identified in male and female p53^R172H^ and male p53^Y217C^ are presented in **Figure 6B**. In all three analyses, mutant p53 was found to localize to the consensus sequence of a unique set of transcription factors, suggesting that the sex specific GOF activity of each mutation may be driven by different sets of transcriptional mediators.

The transcription factor AP-2gamma (*TFAP2C*) is a known regulator of stem cell self-renewal and chemotherapy resistance^51^. We found that in females but not males p53^R172H^ localization is enriched at AP-2gamma DNA-binding motifs. *WNT5A* has been shown to be upregulated in glioma and is associated with poor overall survival^52^. In our model, *Wnt5a* mRNA was significantly upregulated in female p53^R172H^ and down regulated in male p53^R172H^. This difference in gene expression is consistent with greater ChIP-seq signal in females compared to males at the AP-2gamma motif near the TSS of *Wnt5a* (**Figure 6C**). Similarly, p53^Y217C^ localized to the androgen receptor-halfsite motif located upstream of the growth factor Fgf2 in males but not females correlating with sex differences in gene expression (**Figure 6D**). These results suggest that sex differences in mutant p53 GOF gene regulation may be mediated through differential localization to the consensus sequences of known transcription factors upstream of oncogenes.

## Discussion

p53 is the most interrogated protein in cancer research^53^. A PubMed search of p53, TP53, and Trp53 reveals an annual average of 6,000 p53 publications over past five years. Despite this herculean effort, our understanding of the most mutated gene in cancer remains incomplete. An obstacle to fully characterizing WT and mutant p53 activity is its context dependence in function. WT p53 is positioned at the center of a complex network of cellular pathways whose precise activation profile is determined by cell identity and the stimulus activating the p53 response^54^. Similarly, p53 mutations result in a spectrum of cancer phenotypes dictated by the tumor type, co-occurring mutations, and specific p53 mutation^55^. Mounting evidence in both normal development and disease support that sex is a key modulator of p53 function, and that a full understanding of WT and mutant p53 will require incorporating the scope and magnitude of these sex effects.

Most mutations in tumor suppressors prevent gene expression or protein production or result in an unstable protein that is quickly degraded. In contrast, p53 GOF activity selects for stable expression of mutant p53 protein over loss of function mutations or deletions. However, the importance p53 GOF mutations is rarely reflected in *in vitro* and mouse cancer models which more frequently rely on dominant negative p53 constructs, or p53 knockdown or deletion. In GBM, complete p53 loss is an uncommon event. Our analysis of p53 mutant tumors revealed that missense mutations in the DBD are particularly enriched in GBM and CNS cancers compared to other cancers. Together, with previous studies of Li Fraumeni Syndrome demonstrating an association between germline DNA-binding domain mutations and a higher risk of brain tumor development, these findings support a CNS tissue specific advantage of p53 GOF mutation in tumor development^56^. This finding also suggests the fidelity of GBM models could be enhanced by inclusion of clinically-relevant p53 GOF mutations.

In this study we revealed a novel interaction between p53 mutation and cell sex. We found that while p53 deletion increased proliferation, it is insufficient to transform astrocytes and does not induce a sex difference in phenotype. In contrast, introduction of GOF mutant p53 into the same cells was sufficient for transformation, with the magnitude of tumorigenic potential dependent on the mutation and cell sex. The hotspot mutation p53^Y217C^ exhibited the greatest GOF and greatest sex difference of all three tested mutations. Male p53^Y217C^ cells were highly proliferative and tumorigenic while female p53^Y217C^ cells exhibited no GOF phenotype compared to KO. Notably, this relationship is concordant with a male bias in mutation frequency of p53^Y220C^ in humans. p53^R172H^ exhibited the opposite effect with the GOF effect predominant in females with little impact on tumorigenesis in males. In humans this mutation was found in equal prevalence in both sexes. Independent of p53 status, GBM is more common in males than females (incidence ratio of 1.6:1)^57^. If a mutation were to have equal GOF effects in both sexes, we would expect it to present at a similar incidence ratio to GBM overall. A potential explanation for the lack of sex difference in the incidence p53^R175H^ is that the female specific GOF activity compensates for the other factors that would otherwise result in a male bias. The third mutation, p53^Y202C^ had the smallest sex difference in GOF. While males grew faster *in vitro*, both males and females formed tumors *in vivo*. Importantly, these cells were maintained in the same media conditions and *in vivo* growth was observed in both male and female hosts meaning that the sex specific GOF activity must be intrinsic to the sex of the cell and cannot be explained by the influence of acute sex hormones.

Mutant p53 promotes tumorigenesis through the aberrant regulation of oncogenic target genes. Using RNA-seq, we characterized a sex specific and sex independent GOF transcriptional profile – defined as the differential gene expression between mutant p53 and p53 KO – for each missense mutation. All three mutations resulted in sex- and mutation- specific gene expression patterns with thousands of differentially expressed genes. Comparing the GOF transcriptional profiles between sexes revealed three distinct gene expression patterns concordant with the tumorigenic phenotypes. In p53^Y217C^, the male GOF profile was enriched for many cancers pathways, with few pathways enriched in the shared or female specific profiles. This male specific GOF transcriptional profile is consistent with the male specific GOF phenotype. p53^R172H^, which had the greatest female specific GOF phenotype, exhibited enrichment for cancer relevant pathways in both the female specific and shared differentially expressed genes, and no enrichment for cancer pathways in males. This points to a second model where p53^R172H^ acts on a set of female specific and sex independent target genes whose effects interact to drive sex differences in tumor formation. In the third scenario, p53^Y202C^ was only enriched for cancer pathways in the shared component indicating the GOF activity enacted by this mutation is sex independent which is consistent with the similar rates of tumor formation that we observed in these cells. These findings indicate that mutant p53 can have both sex dependent and sex independent GOF activity, and that the impact of each on tumorigenesis is mutation specific.

Lopes-Ramos et al. recently demonstrated that in normal tissues, transcription factors that are not differentially expressed between males and females have different regulatory binding patterns in males and females. The differences in transcription factor binding thus contribute to sex-biased transcriptional programs. In our study, we demonstrated that sex differences in mutant p53 GOF are being driven by differential mutant p53 localization. In both male and female astrocytes, p53^R172H^ localizes to the transcriptional start site of a subset of upregulated genes with greater overall enrichment in females. Whereas in p53^Y217C^, we only observed mutant p53 localization at upregulated genes in males with discernable pattern of p53 binding above input in females. This is consistent with a universal and female biased GOF by p53^R172H^ and a male specific GOF by p53^Y217C^.

Mutant p53 lacks a consensus response element and relies on interactions with other transcription factors for localization. By performing HOMER motif enrichment analysis for regions near the transcriptional start sites of the top upregulated genes, we identified several novel candidate transcription factors that may be contributing to sex differences in mutant p53 localization and gain of aberrant transcriptional output. Understanding the exact mechanisms of sex specific mutant p53 localization and gene expression will require the interrogation of each potential binding partner. However, here we highlight two promising potential co-transactivators: female specific p53^R172H^ enrichment at the AP-2gamma motif upstream of *Wnt5a* and male specific p53^Y217C^ enrichment at the AR-halfsite motif upstream of *Fgf2*. AP-2gamma is an essential transcription factor for several developmental pathways including neural tube development and male gonad development. It also has been shown to act as an oncogene in breast cancer, melanoma, testicular cancer, neuroblastoma, ovarian cancer, and lung cancer ^58–63^. Expression of AP-2gamma is associated with stem cell renewal, proliferation, and therapy resistance^64^. *Wnt5a* is upregulated in GBM and is has been shown to drive proliferation and invasion in cancer stem cell (CSC) populations^65^. Androgen receptor (AR) is known to positively regulate the expression *Fgf2* in prostate cancer^66^. In GBM, FGF2 promotes CSC maintenance through interactions with the receptor FGFR1^67^. AR is expressed in both male and female GBM, and androgen receptor antagonists are being evaluated as novel therapeutic strategies^68^.

As our understanding of cancer heterogeneity has improved, so then has the need for precision medicine to target the specific vulnerabilities within each unique tumor. However, many promising targeted therapies fall short of expectations in clinical trials. Here, we have demonstrated that the same mutations in p53, in the same cell lineage, from otherwise isogenic backgrounds, have unique activity in males and females. Methods for targeting mutant p53 cancers either attempt to directly target mutant p53 with small molecules or exploit vulnerabilities driven by the mutant p53 activity^69^. The influence of cell state on mutant p53 activity suggests that the utility of these therapies may not be universal even across cancers harboring the same p53 mutation. Consequently, understanding the context dependent – including sex - effects of mutant p53 gain-of-function activity is essential to identifying targetable vulnerabilities in mutant p53 cancers.

## Materials and Methods

### Animal Studies Approval

Study was approved in accordance with an animal studies protocol (no. 2018205) approved by the Animal Studies Committee of Washington University School of Medicine per the recommendations of the NIH Guide for the Care and Use of Laboratory Animals.

### Analysis of missense mutations across cancer

TP53 mutation data was aggregated from multiple studies (Supplemental Table S1). The frequency of missense mutations in the DNA binding domain was calculated as a fraction of the total p53 mutations in the data set. Significance was calculated using Fisher’s Exact Test between each cancer type and the pan cancer data set.

### Meta-analysis of mutation incidence in glioblastoma

*TP53* mutation data was collected for 552 male and 442 female patients from The Cancer Genome Atlas (TCGA 2013), the Catalog of Somatic Mutations in Cancer (COSMIC– ALL CNS), and International Agency for Research on Cancer (IARC) datasets. The absolute number of mutations at each codon in the TP53 gene was quantified for each sex independently, and the sex ratio determine by dividing the number of male tumors with a mutation by female tumors with a mutation for each codon. Significant sex differences in mutation frequency were determined for each codon using Fisher’s Exact Test (*p* < 0.05).

### Primary Mouse Astrocyte Isolation and Culture

*Trp53*^*f/-*^ pups were produced by mating male *Trp53*^*-/-*^ mice (B6.129S2-Trp53tm1Tyj/J - The Jackson Laboratory #002101) with *Trp53*^*f/f*^ mice (B6.129P2-Trp53tm1Brn/J - The Jackson Laboratory #008462). Primary astrocytes were isolated from postnatal day 1 pups as previously described^27^. The sex of the pups was determined by PCR of genomic DNA for *Jarid1c/Jarid1d*^*47*^.

Forward: 5’-CTGAAGCTTTTGGCTTTGAG-3’

Reverse: 5’-CCACTGCCAAATTCTTTGG-3’

Astrocytes from at least three male and three female pups were combined for each male and female.

### Immunocytochemistry/Immunofluorescence

Astrocytes were plated on poly-L-lysine (ScienCell) coated coverslips and fixed with 3.2% paraformaldehyde for 10 minutes at room temperature. Fixed cells were immunolabeled for GFAP and p53 (Methods Table 1). Nuclei were counterstained with DAPI and mounted on microscope slides using Immu-Mount (Fisher Scientific). Images were taken using a fluorescent microscope (Olympus Bx60) and processed using Zeis Zen 3.1 software.

### DNA plasmids, cloning, and virus production

Mouse Trp53 transcript variant 1 (NM_011640.3) was cloned into the retroviral vector pMSCV-IRES-GFP II (Addgene plasmid #52107). Trp53 point mutations were introduced using site-directed mutagenesis with the Q5 Site-Directed Mutagenesis Kit (New England Biolabs E0552S). Retrovirus was produced by plasmid transfection into Platinum-E (Plat-E) cells. Lentivirus to express Cre recombinase was produced by transfecting NIH HEK293T cells with Cre-IRES-PuroR plasmid (Addgene plasmid #30205), Δ8.9 and VSV-G. All transfections were performed with FuGENE 6 transfection agent (Promega E2691) in Opti-MEM Reduced Serum Medium (Gibco 31985070) for 5 hours, and the media replaced with standard DMEM/F12, 10% FBS for viral production. Virus was collected 48, 72, and 96 hours post transfection and combined.

### Primary astrocyte genomic alteration

Trp53^mutantp53/-^ astrocytes were produced by transduction with mutant p53-IRES-eGFP retrovirus followed by subsequent transduction with Cre-IRES-Puro lentivirus. Briefly, postnatal day 1 Trp53^f/-^ astrocytes were transduced with a 1:1 ratio of viral media and growth media for 48 hours. Following stable expression of eGFP, cells were sorted by flow-sort for eGFP positive cells. Sorted astrocytes were transduced with a 1:1 ratio of lentiviral-Cre-IRES-puro virus and growth media for 48 hours. Astrocytes were then selected for 1 week in 2.5 ug/ml puromycin (Sigma P8833). Trp53^KO^ astrocytes was achieved by transduction with lentiviral-Cre-IRES-puro virus and selection for 1 week in 2.5 ug/ml puromycin. All primary astrocytes were transfected three passages post isolation.

### Western blot analysis

Total cell lysates were collected in RIPA buffer supplemented with proteinase and phosphatase inhibitors (Roche 11697498001, Roche 04906937001). Protein was separated by electrophoresis using 4- 12% gradient Bis-Tris NuPAGE gels (Invitrogen) and transferred to a nitrocellulose membrane. Membranes were blocked using Odyssey blocking buffer diluted 1:1 in 0.1% PBST. Membranes were then incubated in primary antibodies diluted in blocking buffer overnight at 4°C, followed by secondary antibodies diluted in blocking buffer for 1 hour at room temperature. Blots were imaged using the BioRad ChemiDoc MP Imager and analyzed using the Bio-Rad Image Lab Software V6.1. Primary and secondary antibodies and their corresponding dilution factors can be found in Methods Table 1.

### *In vitro* growth assay

3000 cells were plated in triplicate in 10 ml media in 10 cm tissue culture plates. After incubation in normal tissue culture conditions for 8 days, the cells were washed with 1x phosphate buffered saline and fixed with 100% methanol at room temperature for 6 minutes. Plates were air dried at room temperature. Cells were stained for 1 hour at room temperature with Giemsa stain (Sigma GS500) diluted 1:20 in dH_2_O. Stained plates were washed 3 times in dH_2_O for 5 minutes at room temperature and air dried before imaging using the Imagescanner III (GE Healthcare). Growth was determined by percentage area of the plate covered by cells using ImageJ v1.52a. The global scale was set according to the width of a plate (10 cm), the image flattened, and the color channels split. Using the green channel only, the background was subtracted from each image, and the threshold set to 235 for each image. A circle of equal size was selected with each plate and the total percent area calculated using the “analyze particles” tool. Within each experiment, the percent area for each plate was normalized to the average of the Male KO plates.

### *In vivo* tumorigenesis: Flank implantations

Flank tumors were generated by implanting mutant p53 astrocytes subcutaneously into the left and right-side flanks of NCr nude mice (Taconic). Male and female astrocytes were trypsinized and resuspended in DMEM/F12, and combined at a 1:1 ratio with Matrigel. 100 ul of cell/Matrigel suspension equaling 750,000 male or female cells were injected into opposite flanks of 3 male and 3 female hosts. Flank tumors size was measured weekly using a digital caliper until the largest tumors reached a terminal volume of 1 cm^3^.

### RNA Sequencing

Total RNA was isolated using the RNeasy Mini Kit (Qiagen) from Trp53 KO or Trp53 mutant astrocytes. Samples were prepared according to library kit manufacturer’s protocol, indexed, pooled, and sequenced on an Illumina HiSeq. Basecalls and demultiplexing were performed with Illumina’s bcl2fastq software and a custom python demultiplexing program with a maximum of one mismatch in the indexing read. RNA-seq reads were then aligned to the Ensembl release 76 primary assembly with STAR version 2.5.1a^70^. Gene counts were derived from the number of uniquely aligned unambiguous reads by Subread:featureCount version 1.4.6-p5^71^. Isoform expression of known Ensembl transcripts were estimated with Salmon version 0.8.2^72^. Sequencing performance was assessed for the total number of aligned reads, total number of uniquely aligned reads, and features detected. The ribosomal fraction, known junction saturation, and read distribution over known gene models were quantified with RseQC version 2.6.2^73^.

All gene counts were then imported into the R/Bioconductor package EdgeR^74^ and TMM normalization size factors were calculated to adjust for samples for differences in library size. Ribosomal genes and genes not expressed in the smallest group size minus one samples greater than one count-per-million were excluded from further analysis. The TMM size factors and the matrix of counts were then imported into the R/Bioconductor package Limma^75^. Weighted likelihoods based on the observed mean-variance relationship of every gene and sample were then calculated for all samples with the voomWithQualityWeights^76^. The performance of all genes was assessed with plots of the residual standard deviation of every gene to their average log-count with a robustly fitted trend line of the residuals. Differential expression analysis was then performed to analyze for differences between conditions and the results were filtered for only those genes with Benjamini-Hochberg false-discovery rate adjusted p-values less than or equal to 0.05.

### Sparse PCA analysis

Sparse PCA analysis was performed on normalized gene counts using R/SparsePCA v0.1.2^77^ and variance plotted using the R package plot3D v1.3^78^. Normalized gene counts for all genes with non-zero sparse loadings for the each of the first three principal components were used to generate heatmaps. Heatmaps were generated using R/heatmap3 v1.1.9^79^.

### Pathway analysis

Pathway enrichment analysis was performed on shared and sex specific differentially expressed gene lists using ShinyGO v0.65^80^. Enrichment analysis is based on hypergeometric distribution followed by FDR correction.

### Whole Exome Sequencing

Whole genomic DNA was isolated from 10^6^ cells using the QIAamp DNA Mini Kit (Qiagen 51304). gDNA was fragmented, indexed, and pool. Indexed pools were hybridized with Agilent SureSelectXT Mouse All Exon kit per manufacturer’s instructions and sequenced on an Illumina NovaSeq-6000. Reads were analyzed using a DRAGEN Bio-IT processor (version 0×18101306) running software version 07.021.602.3.8.4. FASTQ files were mapped to mouse reference mm10 and output in CRAM format with duplicates marked. Hard filtered variants in the mouse exome target region were annotated with snpEff v4.3t.

### Chromatin Immunoprecipitation Sequencing

10 million astrocytes were fixed with 1% formaldehyde in DMEM/F12 for 10 minutes are room temperature with gentle rocking. Fixation was quenched by adding glycine to a final concentration of 125 mM. Cells were washed with cold PBS, scraped into 5 ml cold PBS supplemented with protease inhibitors (PI), phosphatase inhibitors (PPI), and PMSF, and transferred to 15 ml conical tubes. Cell suspension were pelleted by centrifugation at 4°C for 5 minutes at 1000 xg and resuspended in 1 ml Farnham lysis buffer (plus PI, PPI, and PMSF) for 10 minutes with gentle vortexing every two minutes to isolate nuclei. Nuclei were pelleted by centrifugation at 4°C, for 5 minutes at 2000 rpm. The nuclear pellet was resuspended in 1 ml RIPA buffer (plus PI, PPI, and PMSF). Sonication was performed on an Epishear probe-in sonicator (Active Motif) at 50% amplitude for 15 cycles of 10 seconds, with 20 seconds of rest between each cycle. Immunopreciptation was performed as previously described, with an antibody recognizing mouse p53 (see Methods Table 1).

Sequencing reads were generated using Illumina NovaSeq S4 (2×150bp) and processed using the ENCODE Transcription Factor and Histone ChIP-Seq processing pipeline v1.1.6 (http://github.com/ENCODE-DCC/chip-seq-pipeline2). The pipeline filtered and mapped reads to the Mus musculus genome (mm10), validated the quality of the data, and generated fold change signal tracks over the inputs using MACS2. Signal tracks of fold enrichment were visualized with the WashU Epigenome browser (https://epigenomegateway.wustl.edu). Motif search for peaks near gene transcript starting sites was conducted using homer v4.8.3 (http://homer.ucsd.edu/homer/index.html).

### Immunohistochemistry of mouse tumors

Mouse flank tumors were dissected, drop fixed in 4% paraformaldehyde overnight at 4°C, and transferred to 30% sucrose. Tumors were imbedded in a paraffin block, sectioned, and mounted on microscope slides. Histological features were determined by hematoxylin and eosin staining. Immunohistochemistry was performed against mouse p53 and Ki67. Antibodies and dilutions are listed in Methods Table 1. Immunohistochemical signal was developed with the ImmPACT DAB Peroxidase Kit (Vector Laboratories, SK-4105).

### Statistics

All *in vitro* experiments were carried out at least three times using cells from three independent thaws and platings. Comparisons between mutation frequencies were determined by Fisher’s Exact Test. Two-tailed Students t-test were used for direct comparisons between mutants in the growth assay and tumor volumes. Differences in tumor growth rate were determined by two-way ANOVA. A *p*-value < 0.05 was considered statistically significant. Correlations between mutational burden and proliferation were calculated using Pearson correlation. Statistics were calculated using GraphPad Prism version 9.0.1.

### Antibodies

**Methods Table 1**

| Antibody | Antigen | Host | Source | Application | Dilution |
| --- | --- | --- | --- | --- | --- |
| 2524 | Mouse p53 | Mouse | Cell Signaling Technology | Western Blot | 1:1000 |
|  |  |  |  | ChIP | 5 µg antibody/25 µg DNA |
|  |  |  |  | ICC | 1:2000 |
| SC-365062 | GAPDH | Mouse | Santa Cruz | Western Blot | 1:200 |
| P53-CM5P-L | P53 | Rabbit | Leica | IHC | 1:200 |
| 13-0300 | GFAP | Rat | Invitrogen | ICC | 1:250 |
| E0431 | Rabbit Immunoglobulin | Swine | Dako | IHC – Secondary/<br>Biotin | 1:750 |
| 926-32212 | Mouse IgG | Donkey | LI-COR | Western Blot –<br>Secondary/<br>IRDye 800CW | 1:50,000 |
| A11077 | Rat IgG | Goat | Invitrogen | ICC – Secondary/<br>Fluor 555 | 1:1000 |

## Supporting information

Rockwell 2021_Supplement

## Acknowledgements

This work was supported by NIH grant RO1 CA174737-06 (JBR), the Children’s Discovery Institute of Washington University and St. Louis Children’s Hospital, and Joshua’s Great Things (JBR). We thank the Alvin J. Siteman Cancer Center at Washington University School of Medicine and Barnes-Jewish Hospital in St. Louis, MO. and the Institute of Clinical and Translational Sciences (ICTS) at Washington University in St. Louis, for the use of the Genome Technology Access Center, which provided RNA-, DNA-, and ChIP-sequencing services. The Siteman Cancer Center is supported in part by an NCI Cancer Center Support Grant #P30 CA091842 and the ICTS is funded by the National Institutes of Health’s NCATS Clinical and Translational Science Award (CTSA) program grant #UL1 TR002345.$We also thank the Alvin J. Siteman Cancer Center at Washington University School of Medicine and Barnes-Jewish Hospital in St. Louis, MO., for the use of the Siteman Flow Cytometry, which provided flow sorting of eGFP positive cells.

