## Supplementary material for "p53 mutations exhibit sex specific gain-of-function activity in gliomagenesis": Rockwell 2021_Supplement

**Supplemental Table 1.** Aggregated studies for p53 mutation analysis.

### Supplemental Table 1

| Cancer group | Studies | Cancer Types | Total number of tumors |
|---|---|---|---|
| **Pan Cancer** | MSK-IMPACT Clinical Sequencing Cohort (MSKCC, Nat Med 2017)[1] | Adenocarcinoma, Adenocarcinoma of Lung, Adenocarcinoma, Mucinous, Breast Neoplasms, Carcinoma, Ductal, Breast, Carcinoma, Hepatocellular, Carcinoma, Lobular, Carcinoma, Non-Small-Cell Lung, Carcinoma, Renal Cell, Carcinoma, Squamous Cell, Carcinosarcoma, Cholangiocarcinoma, Colonic Neoplasms, Colorectal Neoplasms, Cystadenocarcinoma, Serous, DNA Mutational Analysis, Gallbladder Neoplasms, Glioblastoma, Liposarcoma, Lung Neoplasms, Lymphoma, Follicular, Melanoma, Mesothelioma, Neoplasms, Squamous Cell, Neurofibroma, Oligodendroglioma, Osteosarcoma, Ovarian Neoplasms, Pancreatic Neoplasms, Prostatic Neoplasms, Rectal Neoplasms, Sarcoma, Sarcoma, Synovial, Squamous Cell Carcinoma of Head and Neck, Stomach Neoplasms, Thyroid Cancer, Papillary, Urinary Bladder Neoplasms, Uterine Neoplasms, Uveal Neoplasms | 4985 |
| **Breast Cancer** | Breast Cancer (METABRIC, Nature 2012 & Nat Commun 2016)[2,3], Breast Cancer (MSK, Cancer Cell 2018)[4], Breast Cancer (MSK, Nature Cancer 2020)[5], Breast Cancer (MSKCC, NPJ Breast Cancer 2019)[6], Breast Cancer (SMC 2018)[7], Breast Cancer Xenografts (British Columbia, Nature 2015)[8], Breast Fibroepithelial Tumors (Duke-NUS, Nat Genet 2015)[9], Breast Invasive Carcinoma (British Columbia, Nature 2012)[10], Breast Invasive Carcinoma (Broad, Nature 2012)[11], Breast Invasive Carcinoma (TCGA, Cell 2015)[12], Breast Invasive Carcinoma (TCGA, Firehose Legacy), Breast Invasive Carcinoma (TCGA, Nature 2012)[13], Breast Invasive Carcinoma (TCGA, PanCancer Atlas)[14], Metastatic Breast Cancer (INSERM, PLoS Med 2016)[15], The Metastatic Breast Cancer Project (Provisional, February 2020) | Breast, Breast Invasive Cancer, NOS, Breast Invasive Carcinoma, Breast Invasive Carcinoma (NOS), Breast Invasive Ductal Carcinoma, Breast Invasive Mixed Mucinous Carcinoma, Breast Mixed Ductal and Lobular Carcinoma, Ductal Carcinoma In Situ (DCIS), Infiltrating Ductal Carcinoma, Infiltrating Lobular Carcinoma, Invasive Breast Cancer, Invasive Breast Carcinoma, Malignant Phyllodes Tumor of the Breast, Medullary Breast Carcinoma, Metaplastic Breast Cancer, Mixed Carcinoma, Paget Disease of the Nipple | 3219 |

| | | | |
|---|---|---|---|
| **Lung Cancer** | Lung Adenocarcinoma (Broad, Cell 2012)[16], Lung Adenocarcinoma (MSKCC, Science 2015)[17], Lung Adenocarcinoma (OncoSG, Nat Genet 2020)[18], Lung Adenocarcinoma (TCGA, Firehose Legacy), Lung Adenocarcinoma (TCGA, Nature 2014)[19], Lung Adenocarcinoma (TCGA, PanCancer Atlas), Lung Adenocarcinoma (TSP, Nature 2008)[20], Lung Cancer (SMC, Cancer Research 2016)[21], Lung Squamous Cell Carcinoma (TCGA, Firehose Legacy), Lung Squamous Cell Carcinoma (TCGA, Nature 2012), Lung Squamous Cell Carcinoma (TCGA, PanCancer Atlas), Non-Small Cell Cancer (MSKCC, Cancer Discov 2017)[22], Non-small cell lung cancer (MSK, Science 2015)[17], Non-Small Cell Lung Cancer (MSKCC, J Clin Oncol 2018)[23], Non-Small Cell Lung Cancer (TRACERx, NEJM & Nature 2017)[24], Non-Small Cell Lung Cancer (University of Turin, Lung Cancer 2017)[25], Pan-Lung Cancer (TCGA, Nat Genet 2016)[26], Small Cell Lung Cancer (CLCGP, Nat Genet 2012)[27], Small Cell Lung Cancer (Johns Hopkins, Nat Genet 2012)[28], Small Cell Lung Cancer (U Cologne, Nature 2015)[29], Small-Cell Lung Cancer (Multi-Institute, Cancer Cell 2017)[30], Thoracic PDX (MSK, Provisional) | Combined Small Cell Lung Carcinoma, Large Cell Lung Carcinoma, Large Cell Neuroendocrine Carcinoma, Lung Adenocarcinoma, Lung Adenosquamous Carcinoma, Lung Neuroendocrine Tumor, Lung Squamous Cell Carcinoma, Non-Small Cell Lung Cancer, Pleomorphic Carcinoma of the Lung, Pleural Mesothelioma, Poorly Differentiated Non-Small Cell Lung Cancer, Sarcomatoid Carcinoma of the Lung, Small Cell Lung Cancer | 3727 |
| **Skin** | Acral Melanoma (TGEN, Genome Res 2017)[31], Desmoplastic Melanoma (Broad Institute, Nat Genet 2015)[32], Melanoma (Broad/Dana Farber, Nature 2012)[33], Melanoma (MSKCC, NEJM 2014)[34], Melanomas (TCGA, Cell 2015)[35], Metastatic Melanoma (DFCI, Nature Medicine 2019)[36], Metastatic Melanoma (DFCI, Science 2015)[37], Metastatic Melanoma (MSKCC, JCO Precis Oncol 2017)[38], Metastatic Melanoma (UCLA, Cell 2016)[39], Skin Cutaneous Melanoma (Broad, Cell 2012)[40], Skin Cutaneous Melanoma (TCGA, Firehose Legacy), Skin Cutaneous Melanoma (TCGA, PanCancer Atlas)[14], Skin Cutaneous Melanoma (Yale, Nat Genet 2012)[41], Skin Cutaneous Melanoma(Broad, Cancer Discov 2014)[42] | Acral Melanoma, Cutaneous Melanoma, Desmoplastic Melanoma, Melanoma, Melanoma of Unknown Primary | 257 |

| | | | |
|---|---|---|---|
| **Prostate** | Prostate Adenocarcinoma (Broad/Cornell, Cell 2013)[43], Prostate Adenocarcinoma (Broad/Cornell, Nat Genet 2012)[44], Prostate Adenocarcinoma (CPC-GENE, Nature 2017)[45], Prostate Adenocarcinoma (Fred Hutchinson CRC, Nat Med 2016)[46], Prostate Adenocarcinoma (MSK, Eur Urol 2020)[47], Prostate Adenocarcinoma (MSKCC, Cancer Cell 2010)[48], Prostate Adenocarcinoma (MSKCC/DFCI, Nature Genetics 2018)[49], Prostate Adenocarcinoma (SMMU, Eur Urol 2017)[50], Prostate Adenocarcinoma (TCGA, Firehose Legacy), Prostate Adenocarcinoma Organoids (MSKCC, Cell 2014)[51], Prostate Cancer (MSKCC, JCO Precis Oncol 2017)[52] | Prostate Adenocarcinoma, Prostate Neuroendocrine Carcinoma, Prostate Small Cell Carcinoma | 970 |
| **Colon** | Colon Adenocarcinoma (CaseCCC, PNAS 2015)[53], Colon Cancer (CPTAC-2 Prospective, Cell 2019)[54], Colorectal Adenocarcinoma (DFCI, Cell Reports 2016)[55], Colorectal Adenocarcinoma (Genentech, Nature 2012)[56], Colorectal Adenocarcinoma (TCGA, Firehose Legacy), Colorectal Adenocarcinoma (TCGA, PanCancer Atlas)[14], Colorectal Adenocarcinoma Triplets (MSKCC, Genome Biol 2014)[57], Metastatic Colorectal Cancer (MSKCC, Cancer Cell 2018)[58], Rectal Cancer (MSK,Nature Medicine 2019)[59] | Colorectal Adenocarcinoma, Colon Adenocarcinoma, Rectal Adenocarcinoma, Mucinous Adenocarcinoma | 2342 |
| **CNS** | Anaplastic Oligodendroglioma and Anaplastic Oligoastrocytoma (MSKCC, Neuro Oncol 2017)[60], Brain Lower Grade Glioma (TCGA, Firehose Legacy), Brain Tumor PDXs (Mayo Clinic, 2019), Glioblastoma (Columbia, Nat Med. 2019)[61], Glioblastoma (TCGA, Cell 2013)[62], Glioblastoma (TCGA, Nature 2008)[63], Glioblastoma Multiforme (TCGA, Firehose Legacy), Glioma (MSK, Nature 2019)[64], Glioma (MSKCC, Clin Cancer Res 2019)[65], Low-Grade Gliomas (UCSF, Science 2014)[66], Merged Cohort of LGG and GBM (TCGA, Cell 2016)[67], Pheochromocytoma and Paraganglioma (TCGA, Firehose Legacy) | Anaplastic Astrocytoma, Anaplastic Oligoastrocytoma, Anaplastic Oligodendroglioma, Astrocytoma, Desmoplastic/Nodular Medulloblastoma, Diffuse Astrocytoma, Diffuse Glioma, Glioblastoma, Glioblastoma Multiforme, Gliosarcoma, High-Grade Glioma, NOS, Large Cell/Anaplastic Medulloblastoma, Low-Grade Glioma (NOS), Medulloblastoma, Oligoastrocytoma, Oligodendroglioma, Pheochromocytoma, Rosette-forming Glioneuronal Tumor of the Fourth Ventricle | 1979 |
| **Glioblastoma** | Glioblastoma (Columbia, Nat Med. 2019)[68], Glioblastoma (TCGA, Cell 2013)[62], Glioblastoma Multiforme (TCGA, Firehose Legacy) | Glioblastoma Multiforme | 316 |

### Supplementary Table 1 References

23. Rizvi H, Sanchez-Vega F, La K, et al. Molecular determinants of response to anti-programmed cell death (PD)-1 and anti-programmed death-ligand 1 (PD-L1) blockade in patients with non-small-cell lung cancer profiled with targeted next-

generation sequencing. *J Clin Oncol*. 2018;36(7):633-641. doi:10.1200/JCO.2017.75.3384

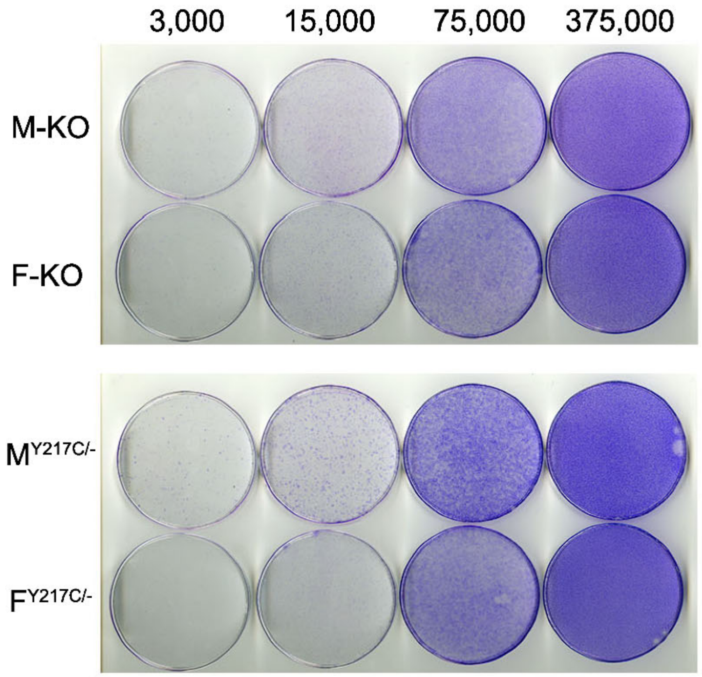


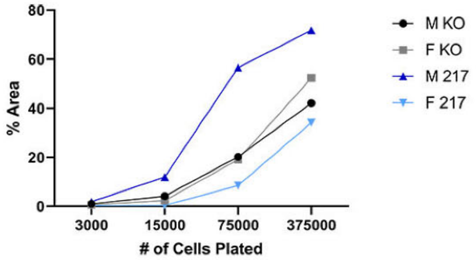


**Supplemental Figure 1.** Titration of p53$^{Y217C}$ foci assay. **A.** Male and female p53 KO and p53$^{Y217C}$ astrocytes were plated at a 5-fold dilution in 10 cm plates and incubated for five days before fixing and staining and with Giemsa stain. **B.** Quantification of foci assay as measured by % area covered by stained nuclei.

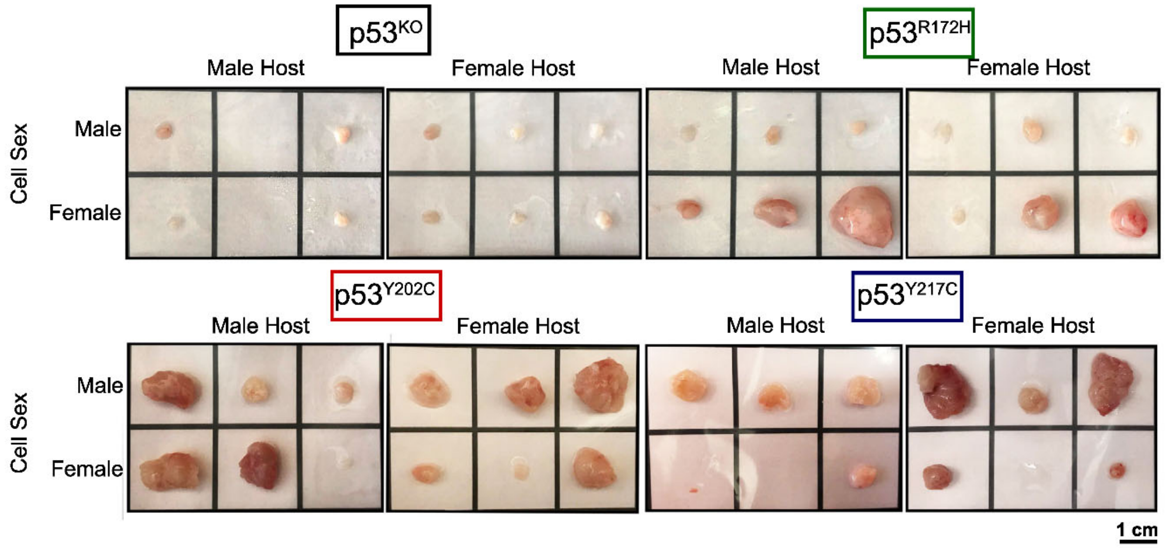


**Supplemental Figure 2.** Images of harvested flank tumors or recovered Matrigel pellet from mice injected with male and female p53 KO (black), p53$^{R172H}$ (green), p53$^{Y202C}$ (red), and p53$^{Y217C}$ (blue). Where blank squares are present, no tumor or Matrigel pellet was recovered.

A.

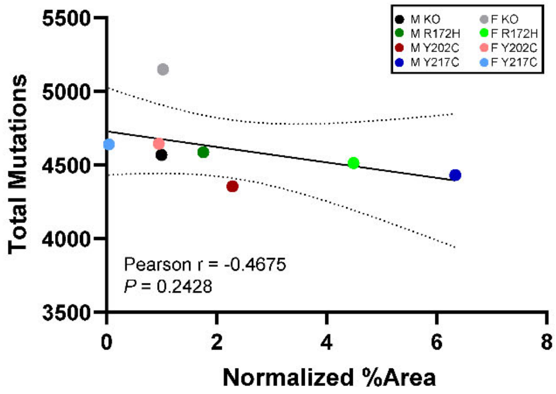


B.

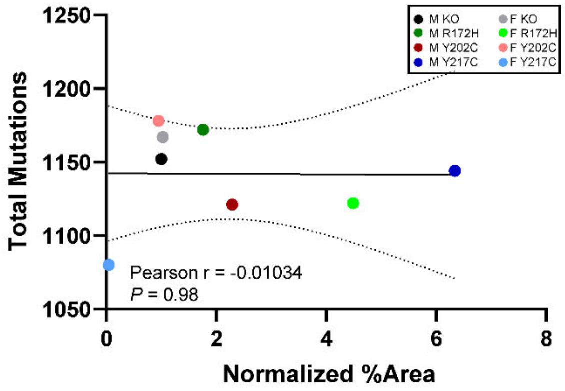


**Supplemental Figure 3.** Cell growth phenotype does not correlate with mutation burden. Pearson correlation of the relationship between growth as measure by normalized percent area from the foci assay and the total number of mutations (**A**) or the number of missense or nonsense mutations (**B**) identified in whole exome sequencing.

**Supplemental Table 2.** All identified mutant genes in male or female p53 KO, p53$^{R172H}$, p53$^{Y202C}$, and p53$^{Y217C}$ astrocytes. A (+) indicates a homologous mutation in the given gene in the corresponding mutation p53 cell line.

| | Male | | | | Female | | | |
|---|---|---|---|---|---|---|---|---|
| **Gene Name** | **KO** | **R172H** | **Y202C** | **Y217C** | **KO** | **R172H** | **Y202C** | **Y217C** |
| 1600014C10Rik | | + | + | + | + | + | + | + |
| 1810065E05Rik | | | + | + | | | | + |
| 2210407C18Rik | | | + | + | | | | + |
| 2810021J22Rik | | | + | + | | | | + |
| 4933427D14Rik | + | + | + | + | + | + | + | + |
| 6330403K07Rik | + | + | + | + | + | + | + | + |
| 6330408A02Rik | | + | + | + | + | + | + | + |
| Akap10 | | | + | + | + | + | | + |
| Alox8 | + | + | + | + | | + | + | + |
| Aloxe3 | + | + | + | + | + | + | + | + |
| Arap2 | + | + | + | + | + | + | + | + |
| Aspa | + | + | + | + | + | + | + | + |
| Aurkb | + | + | + | + | + | + | + | + |
| Auts2 | | + | + | + | + | + | + | + |
| AV320801 | + | + | + | + | | | | |
| Bean1 | | + | + | + | + | + | + | + |
| Btnl10 | | | + | + | | | | + |
| Calcoco2 | | | | | | + | | |
| Ccdc92b | + | + | + | + | + | + | + | + |
| Chd3 | + | + | + | + | + | + | + | + |
| Cntrob | + | + | + | + | + | + | + | + |
| Cyb5d1 | + | + | + | + | + | + | + | + |
| Dhtkd1 | + | + | + | + | + | + | + | + |
| Dmbt1 | + | + | + | + | + | + | + | + |
| Dnah9 | | | + | + | | | | + |
| Dux | + | + | + | + | + | + | + | + |
| Elac2 | | | + | | | + | | + |
| Fam114a2 | | | + | + | | + | | + |
| Fam64a | + | + | + | + | + | + | + | + |
| Fam83g | | | + | + | | + | | + |
| Fat2 | | | + | + | | | | + |
| Fbxw10 | | | + | + | | + | | + |
| Galnt10 | | | + | | | | | + |
| Gemin5 | | | + | + | | | | + |
| Glod4 | + | + | + | + | + | + | + | + |
| Glp2r | + | + | + | + | + | + | + | + |
| Gm12250 | | | + | + | | + | | + |
| Gm12253 | | | + | + | | + | | + |
| Gsg2 | + | + | + | + | + | + | + | + |
| Gucy2e | + | + | + | + | + | | + | + |
| Hebp1 | | | + | | | | | |
| Hes7 | + | + | + | + | + | + | + | + |
| Hic1 | + | + | + | + | + | + | + | + |
| Hoxa13 | + | + | + | + | + | + | + | + |

| | | | | | | | | |
|---|---|---|---|---|---|---|---|---|
| Iba57 | | | + | + | | | | + |
| Ifi30 | | + | + | + | + | + | + | + |
| Igtp | | | + | + | | | | + |
| Irgm2 | | | + | + | | + | | + |
| Itgae | + | + | + | + | + | + | + | + |
| Kif1c | + | + | + | + | + | + | + | + |
| Kndc1 | | + | + | + | + | + | + | + |
| Larp1 | | | + | + | | + | | + |
| Lig1 | | + | + | + | + | + | + | + |
| Lypd8 | | | + | + | | + | | + |
| Mfap3 | | | + | + | | + | | + |
| Mmp1a | | + | + | + | + | + | + | + |
| Mroh2a | + | + | + | + | + | + | + | + |
| Mrpl22 | | | + | + | | | | + |
| Mrpl55 | | | + | + | | | | + |
| Muc4 | + | + | + | + | + | + | + | + |
| Muc6 | + | + | + | + | + | + | + | + |
| Myh1 | | | + | + | | | | + |
| Myh8 | | | + | | | | | + |
| Myocd | | | + | + | | | | + |
| Nadk2 | | + | + | + | + | + | + | + |
| Nlgn2 | + | + | + | + | + | + | + | + |
| Nlrp1a | + | + | + | + | + | + | + | + |
| Nlrp1b | + | + | + | + | + | + | + | + |
| Nmur2 | | | + | + | | + | | + |
| Obscn | | | + | + | | + | | + |
| Olfr224 | | | + | + | | + | | + |
| Olfr311 | | | + | + | | + | | + |
| Olfr313 | | | + | + | | | | + |
| Olfr314 | | | + | + | | + | | + |
| Olfr316 | | | + | + | | + | | + |
| Olfr317 | | | + | + | | | | + |
| Olfr319 | | | + | + | | + | | + |
| Olfr320 | | | + | + | | | | + |
| Olfr322 | | | + | + | | + | | + |
| Olfr323 | | | + | + | | | | + |
| Olfr324 | | | + | + | | + | | + |
| Olfr325 | | | + | + | | | | + |
| Olfr328 | | | + | + | | | | + |
| Olfr329-ps | | | + | + | | | | + |
| Olfr331 | | | + | + | | | | + |
| Olfr332 | | | + | + | | + | | + |
| Otop1 | + | + | + | + | + | + | + | + |
| Ovca2 | + | + | + | + | + | + | + | + |
| Pitpnm3 | + | + | + | + | + | + | + | + |
| Prpsap2 | | | + | + | | | | + |
| Ptpmt1 | | + | + | + | + | + | + | + |

| | | | | | | | | |
|---|---|---|---|---|---|---|---|---|
| Rai1 | | | + | + | | | | + |
| Rbmy | | | + | | | | | |
| Rnf167 | | + | + | + | + | + | + | + |
| Rnmtl1 | + | + | + | + | + | + | + | + |
| Ror2 | + | + | + | + | + | + | + | + |
| Rpa1 | + | + | + | + | + | + | + | + |
| Sema3a | + | + | + | + | + | + | + | + |
| Sfi1 | + | + | + | + | + | + | + | + |
| Sftpb | | + | + | + | + | + | + | + |
| Slc6a20b | + | + | + | + | + | + | + | + |
| Slc6a4 | + | + | + | + | + | + | + | + |
| Smcr8 | | | + | + | | | | + |
| Smg6 | + | + | + | + | + | + | + | + |
| Spns3 | + | + | + | + | + | + | + | + |
| Srebf1 | | | + | + | | | | + |
| Tekt1 | + | + | + | + | + | + | + | + |
| Thap1 | | | + | | | | | |
| Timm22 | + | + | + | + | + | + | + | + |
| Tmem102 | + | + | + | + | + | + | + | + |
| Tmem134 | | + | + | + | + | + | + | + |
| Tmem267 | | + | | | + | | | |
| Tom1l2 | | | + | + | | | | + |
| Top3a | | | + | + | | | | + |
| Trerf1 | | + | + | + | + | + | + | + |
| Trim16 | | | + | + | | | | + |
| Trim17 | | | + | + | | + | | + |
| Trim58 | | | + | + | | | | + |
| Trp53 | | + | + | + | | + | + | + |
| Trpv1 | + | + | + | + | + | + | + | + |
| Trpv3 | + | + | + | + | + | + | + | + |
| Tvp23b | | | + | + | | | | + |
| Ugt1a1 | + | + | + | + | + | + | + | + |
| Usp43 | + | + | + | + | + | + | + | + |
| Vezf1 | | | | | | + | | + |
| Xaf1 | + | + | + | + | + | + | + | + |
| Zc3h7a | | + | + | + | + | + | + | + |
| Zfp286 | | | + | + | | | | + |
| Zfp287 | | | + | + | | | | + |
| Zfp39 | | | + | + | | | | + |
| Zfp692 | | | + | + | | | | + |
| Zfp991 | + | | + | + | + | + | + | |
| Zzef1 | + | + | + | + | + | + | + | + |

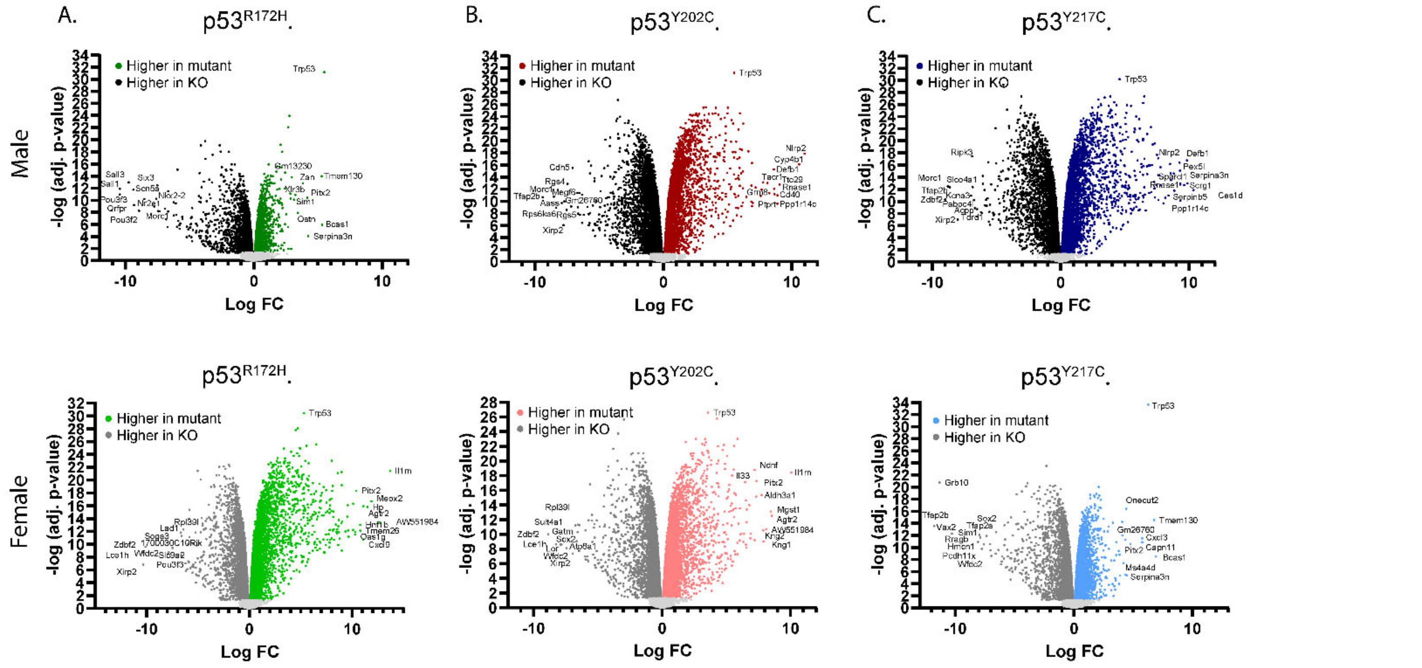


**Supplemental Figure 4.** Volcano plots displaying all significant differential gene expression (adjusted p-value < 0.05) between mutant p53 and p53 KO astrocytes within each sex for **A.** p53$^{R172H}$ (green), **B.** p53$^{Y202C}$ (red), **C.** p53$^{Y217C}$ (blue). The top ten upregulated and downregulated genes by logFC and *Trp53* are labeled in each plot. Genes below the significance threshold are colored light grey.
